# The Juvenile Idiopathic Arthritis (JIA) Susceptibility Locus, *IL2RA*, Includes An Intronic Enhancer That Is Attenuated By JIA-Associated Genetic Variants

**DOI:** 10.1101/554006

**Authors:** Kaiyu Jiang, Yungki Park, Evan Tarbell, Tao Liu, James N. Jarvis

## Abstract

Like most rheumatic diseases, genetic risk loci for juvenile idiopathic arthritis (JIA) are highly enriched for H3K4me1/H3K27ac histone marks, epigenetic signatures often associated with enhancer function. We sought to determine whether an H3K4me1/H3K27ac region of the JIA risk locus, *IL2RA* exhibits enhancer function and whether enhancer function is altered by JIA-associated genetic variants. We used a conventional luciferase reporter assay to query enhancer function across an H3K4me1/H3K27ac-marked region within the first intron of the *IL2RA* gene. We used the same approach to query the effects of 4 single nucleotide polymorphisms (SNPs) on enhancer function. We identified a 657 base pair region within the first intron of *IL2RA* that demonstrated brisk enhancer function. Enhancer activity was observable in both Jurkat T cells and myeloid HL60 cells. We identified 2 genetic variants, rs117119468 (C->T), and rs12722502 (C->T), that attenuated enhancer activity in this 657 bp region. *Conclusions* – The JIA-associated risk locus, *IL2RA*, contains at least one functional enhancer that is active in both lymphoid and myeloid cells. We identified 2 genetic variants that attenuate activity of this enhancer, making them strong candidates as causal variants.

**Author Summary:** One of the challenges facing the field of genetics as applied to complex traits is the fact that many of the risk-associated genetic variants are located in non-coding regions of the genome. Thus, clarifying the mechanisms through which genetic variants contribute to disease risk requires consideration of the non-coding functions of the risk regions. We and others have shown that many of these risk regions include enhancers, DNA elements that act like rheostats to fine-tune gene expression to fit specific physiologic situations. In this paper, we demonstrate that the juvenile idiopathic arthritis (JIA)- associated genetic risk locus, *IL2RA*, includes an intronic enhancer that is active in cells of both the adaptive and innate immune systems (i.e., the parts of the immune system that require “immune memory” and the parts that don’t). We also identified 2 genetic variants that reduce the function of this enhancer. This is the first discovery of so called “causal variants” that affect non-coding functions in this common childhood disease.

## Introduction

Juvenile idiopathic arthritis (JIA) is a term used to describe a heterogeneous group of childhood-onset illnesses that share the common feature of chronic hypertrophy and inflammation of synovial membranes. JIA represents a classic “complex trait,” in which twin and family studies support a genetic component [1, 2], but where risk conferred by any single disease-associated genetic locus is small.

Genome-wide association studies (GWAS) [2, 3] and genetic fine mapping studies [4] have provided us with a treasure trove of information regarding genetic risk for JIA. The next important tasks for the field are to identify the actual genetic variants that confer risk, identify the cells in which they act, and, finally, identify the specific cellular processes altered by those variants [5]. In accomplishing these tasks, we face several challenges: (1) genetic risk loci are scattered throughout the genome [6, 7] and likely include variants not identified in GWAS or genetic fine mapping studies; and (2) the risk loci consist of large haplotypes that contain multiple genetic variants and functional elements other than genes. Our group has demonstrated, for example, that the JIA risk loci are highly enriched (compared with genome background) for H3K4me1/H3K27ac histone marks, epigenetic signatures often associated with enhancer function [8, 9]. Another serious challenge we face in developing a mechanistic understanding of genetic risk in JIA is related to the large number of known genetic variants (>18,000) within the established risk haplotypes. At the present time, the field has no established method for screening these variants to identify the most promising candidates to test in functional assays. However, the chromatin features within a risk locus can provide inferences regarding the genetic/genomic function of the risk region as well as reduce the number of candidate variants to be tested in functional assays.

In this paper, we report on non-coding functions within the *IL2RA* locus. *IL2RA* was identified as a JIA risk locus using both candidate gene [10] and genetic fine mapping [4] approaches. The linkage disequilibrium (LD) block that includes the SNPs used to identify this locus encompasses the region from chr10:6,071,347-6,097,283 (hg19 assembly) and consists of a large region within the first intron of the *IL2RA* (CD25) gene, as shown in **Figure 1**. The gene, *IL2RA*, encodes for the alpha chain of the IL2 receptor. Because of the importance of IL2 in a broad spectrum of adaptive immune functions as well as particular interest in CD25+FoxP3+ regulatory T cells in JIA [11], it has generally been assumed that variants in this locus must impact *IL2RA* expression and adaptive immune function. However, in gene expression studies of peripheral blood mononuclear cells [12], whole blood [13], neutrophils [14], and CD4+ T cells [15], we have never identified *IL2RA* as a differentially expressed gene when we compare children with active polyarticular JIA to healthy children. We therefore sought to determine whether this JIA risk region might also possess non-coding functions. In addition, we sought to determine the effects of JIA-associated genetic variants on non-coding functions within the *IL2RA* risk locus.

**Figure 1–.**
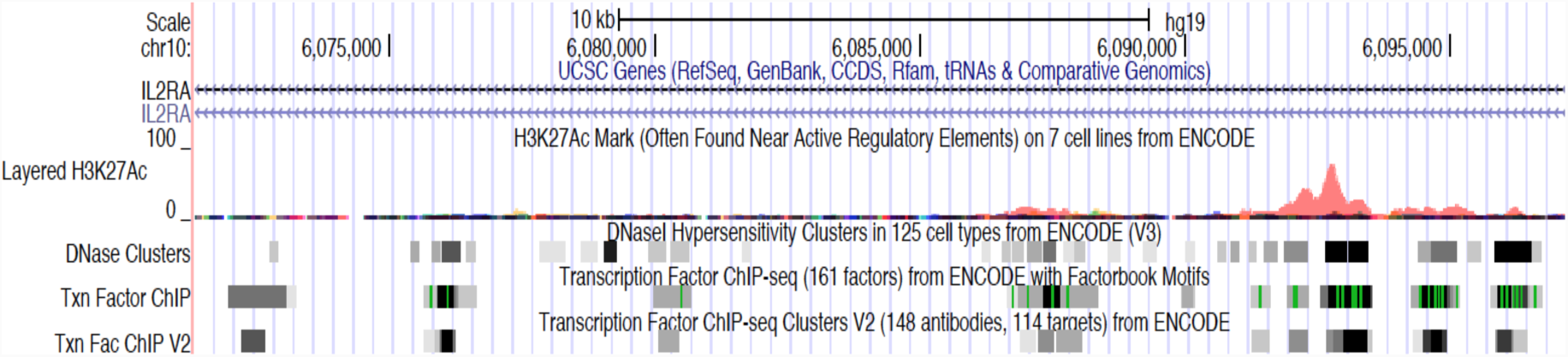
Genome browser screen shot showing the entire *IL2RA* genetic risk locus defined by the SNP, rs7909519. The risk locus encompasses most of the first intron of the *ILRA* gene, spanning chr10:6,071347-6,097,283 (hg19). ENCODE DNase1 hypersensitivity and transcription factor (TF) binding sites are shown on the lower tracks. An H3K27ac ChIPseq peak (orange peak) is identified in the first intron and co-localizes with dense TF binding. H3K4me1 peaks also co-localize to this area (data not shown). ChIPseq data from human neutrophils has also identified H3K4me1/H3K27ac peaks in this region[8].

**Figure 2–.**
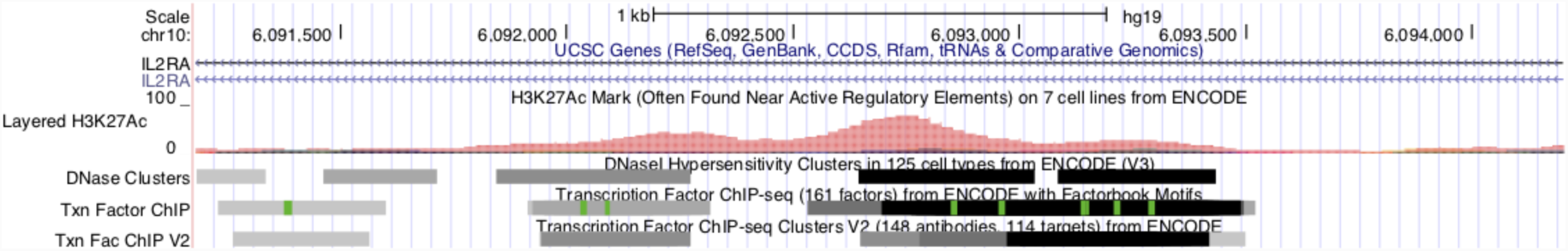
UCSC genome browser view of a select region within the *IL2RA* haplotype within the first intron of the *IL2RA* gene. This region is characterized by dense TF binding and H3K27ac histone peaks (ENCODE and *Roadmap Epigenomics* data). These chromatin features in this region are identical in both human CD4+ T cells and neutrophils.

## Results

### The IL2RA locus includes an intronic enhancer that is active in both lymphoid and myeloid cells

We used a reporter assay to assess enhancer function within the selected 657 bp region as described in the *Methods* section. We tiled two constructs (533bp and 539bp) across the region of interest and tested each construct for luciferase production. We identified the two constructs (chr10:6092661-6093199 and chr10:6092785-6093317) with brisk enhancer activity that was observable in Jurkat T cells as well as myeloid HL60 cells. These results are summarized in **Figure 3**. Note that the region (chr10:6,089,866-6,090,267) that is nearest the tag SNP use to identify *IL2RA* as a susceptibility locus, rs7909519 (chr10:6,089,841), does not show constitutive enhancer function, even though the construct we used has the common allele, *not* rs7909519.

**Figure 3–.**
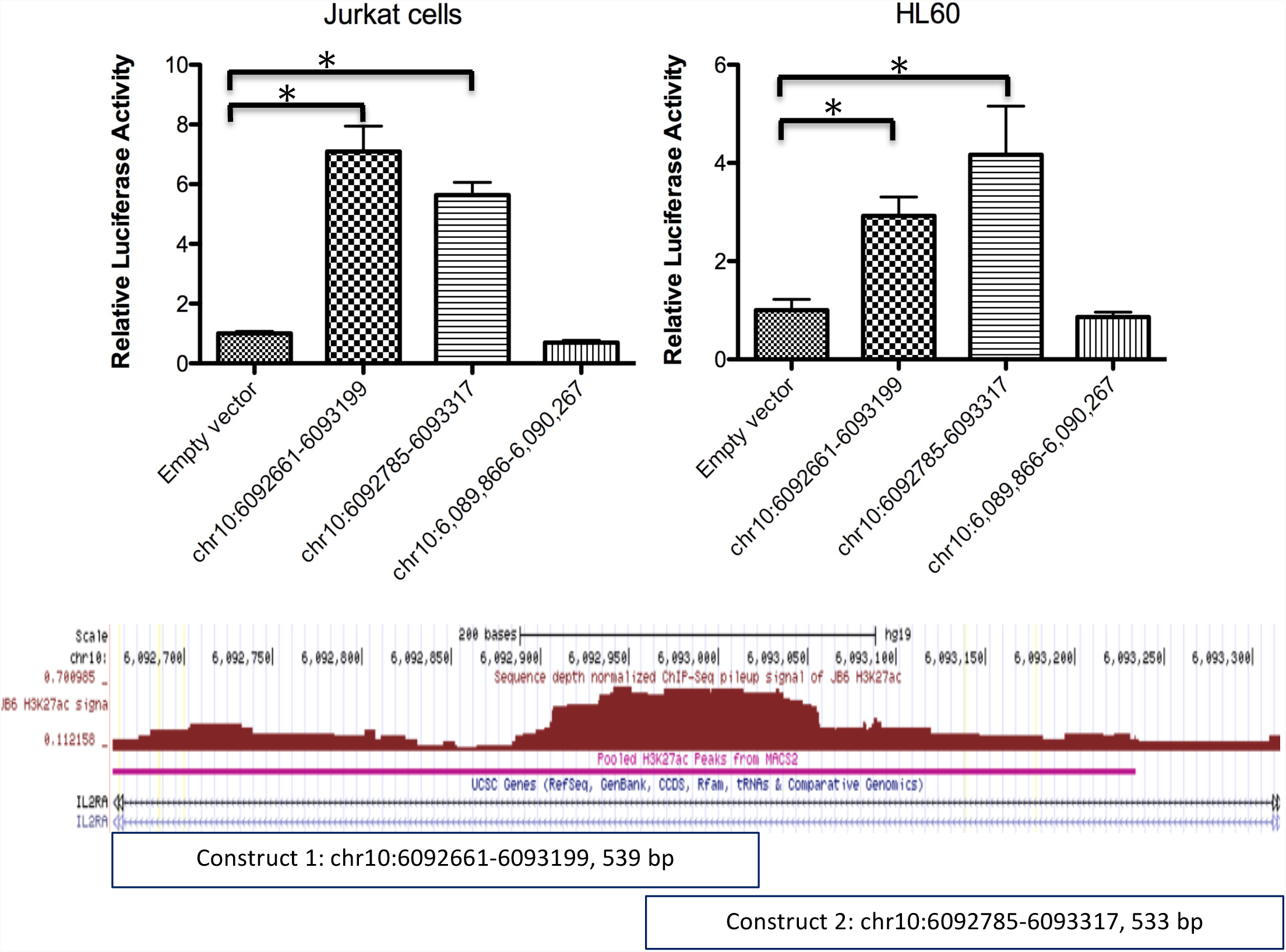
Bar graphs summarizing results from luciferase assays assessing enhancer activity at the *IL2RA* locus in Jurkat T cells (**A**) and HL60 cells (**B**). Each bar represents results from 539bp and 533 constructs, respectively, with the constructs tiled across an H3K4me1/H3K27ac-marked region within JIA-associated haplotype in the first intron of the *IL2RA* gene. Chromosome coordinates for each construct are indicated beneath each bar. The negative control consisted of cells transfected with the blank pGL4.23 vector (which contains the SV40 promoter but is inefficient at driving luciferase expression). Enhancer activity, which increased luciferase expression in Jurkat T cells and in HL-60 cells, is detected within this region. Note that the bar on the far right shows results from the region that includes the index SNP rs7909519. This region does not show enhancer activity. **C**. University of California Santa Cruz (UCSC) Genome Browser screen shot showing the specific region queried by the reporter constructs. The region H3K27ac marked by histone marks is indicated by the brown peak (Roadmap Epigenomics data) and horizontal magenta line (our neutrophil data, GSE66896).

### Genetic variants identified from whole genome sequencing (WGS) of children with JIA alter enhancer function within the IL2RA locus

We next sought to test the effects of genetic variants within the identified enhancer. For these experiments, we first tested 2 variants, rs117119468 (C->T), and rs12722502 (C->T), that we had identified on whole genome sequencing (WGS)[7] of a small number of children with polyarticular JIA. For these experiments, we generated one construct across the region with demonstrated enhancer function (chr10:6092661-6093317, 657bp). The construct was used as template to generate genetic variants using site-directed mutagenesis as described in the *Methods* section. One variant, rs117119468, completely abolished enhancer function within this locus in Jurkat T cells and in HL60 cells. Another, rs12722502, reduced function by 30% in Jurkat but had no effect in HL60 cells. These results are shown in **Figure 4**.

**Figure 4–.**
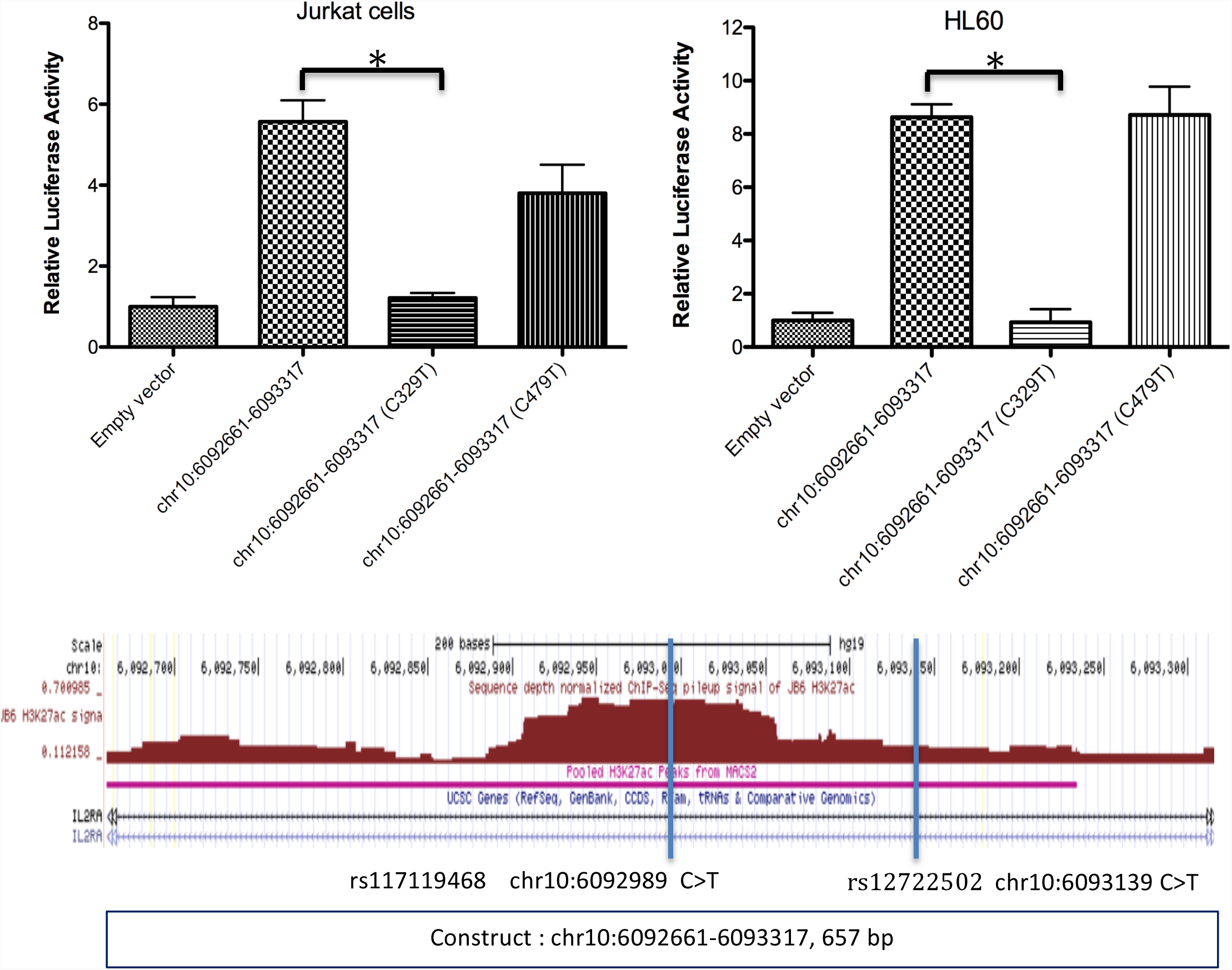
Bar graph showing the effects of JIA-associated genetic variants on enhancer activity in Jurkat T cells (**A**) and in HL60 cells (**B**). The first bar represents the same negative control as shown in *Figure 3*. The next bar represents the region queried in *Figure* 3 using a construct containing the common allele. The third bar shows results using a construct containing the rare variant, chr10:6092989 C->T, identified on WGS of children with polyarticular JIA [10]. This variant almost completely abolishes enhancer function in Jurkat T cells and in HL60 cells. A second variant, chr10:6093139 C->T, reduced luciferase production in Jurkat cells but not in HL60 cells (fourth bar). The fifth bar, on the far right, once again shows that the region that includes the index SNP does not actually have functional activity. **C**. University of California Santa Cruz (UCSC) Genome Browser screen shot showing the specific region queried by the reporter constructs. The vertical blue lines represent the positions of the chr10:6092989 C->T and chr10:6093139 C->T SNPs queried in these assays. The region marked by H3K27ac histone marks is indicated by the brown peak (Roadmap Epigenomics data) and horizontal magenta line (our neutrophil data, GSE66896).

We next tested two common genetic variants annotated in the 1000 Genomes Project data (rs370928127 A>G and rs552847047 G>T). Neither of these variants had any affect on enhancer activity within this region (**Figure 5**). These experiments demonstrate unequivocally that the *IL2RA* locus has enhancer function and that function is impeded by JIA-associated genetic variants.

**Figure 5–.**
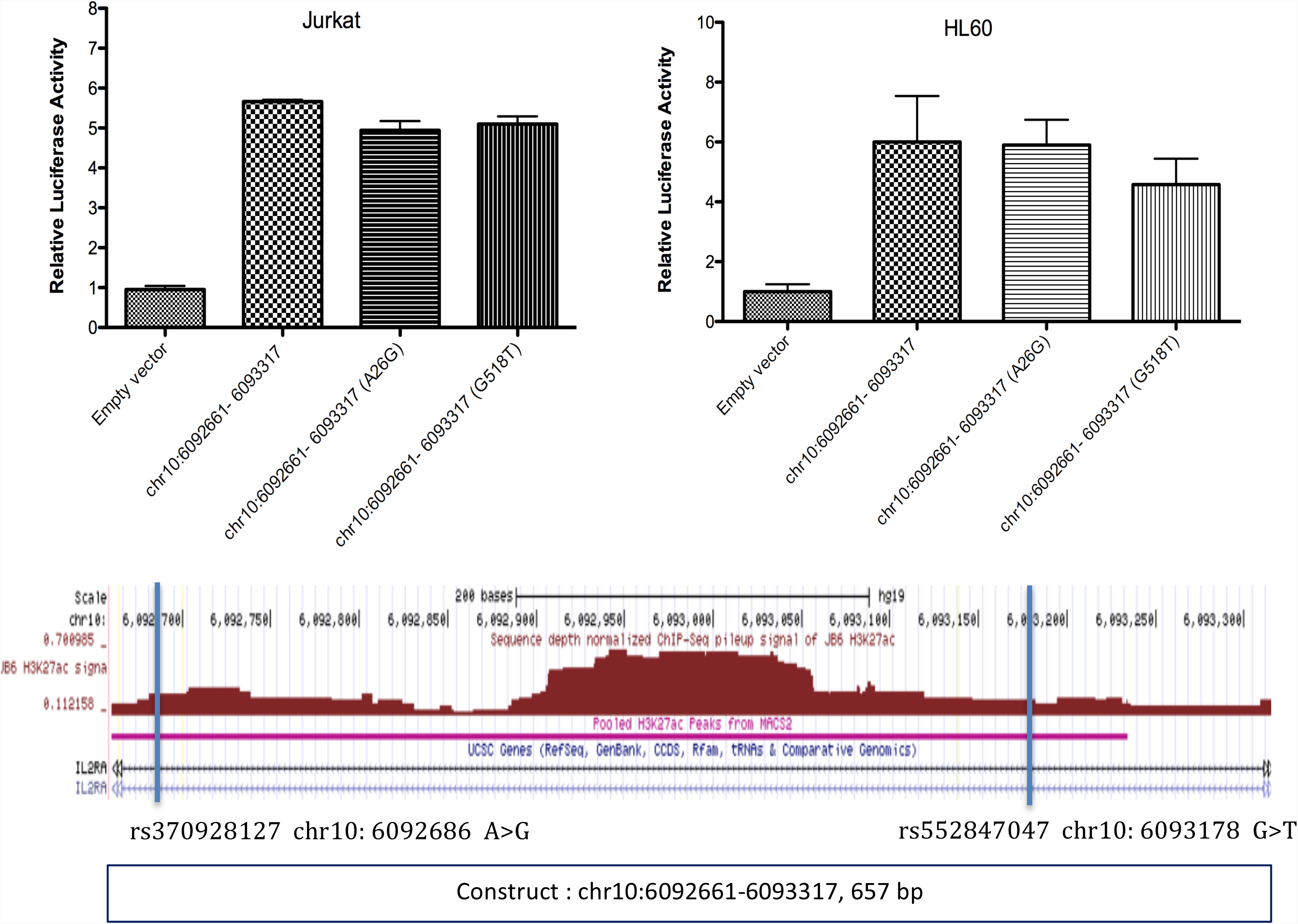
– Bar graph showing the effects of two additional variants annotated in the 1000 Genomes Project data, rs370928127 A->G and, rs552847047 G->T, on enhancer activity in Jurkat T cells (**A**) and in HL60 cells (**B**). The first bar represents the same negative control as above. The next bar represents the region queried above using a construct containing the common allele. The third and forth bar shows neither of these variants had any affect on enhancer activity within this region. **C.** University of California Santa Cruz (UCSC) Genome Browser screen shot showing the specific region queried by the reporter constructs. The vertical blue lines represent the positions of the two variants, rs370928127 A->G and, rs552847047 G->T, queried in these assays. The region marked by H3K27ac histone marks is indicated by the brown peak (Roadmap Epigenomics data) and horizontal magenta line (our neutrophil data, GSE66896).

## Discussion

One of the limitations of GWAS studies is the fact that the index SNPs used to identify risk regions aren’t necessarily those that exert the biological effects that confer disease risk. This is because the index SNPs used to identify risk loci are in linkage disequilibrium with dozens, hundreds, or thousands of other SNPs, any of which may be the one(s) exerting the biological effects that lead to disease risk. The JIA haplotypes therefore present a challenge, as they contain many features including but not limited to genes. We have reported, for example, that the JIA risk loci are highly enriched (compared to genome background) for H3K4me1/H3K27ac histone marks, epigenetic signatures typically associated with enhancer function [8, 9]. JIA is not unique in this regard; the risk loci for most autoimmune diseases show enrichment for epigenetic signatures of enhancer function[16]. The reverse is also true: if one maps enhancer elements in specific cell types, the mapped regions are highly enriched in GWAS-identified SNPs [17] for diseases that affect those cells/tissues. However, chromatin signatures do not, by themselves, establish unequivocally the presence of an active enhancer in a specific region; enhancer function must be determined empirically. In this paper, we used a conventional reporter assay to assess enhancer function across a specific region of the *IL2RA* haplotype block that was characterized by typical chromatin features of enhancer function: open chromatin, abundant transcription factor binding (ENCODE and Roadmap Epigenomics data), and flanking H3K4me1/H3K27ac histone marks. We identified a 657 bp region with strong enhancer function. Furthermore, this enhancer was active in both lymphoid (Jurkat T cells) and myeloid (HL60) cells. The finding that this enhancer operates in both lymphoid and myeloid cells was not surprising. Hematopoetic cells also express common as well as cell-type specific enhancers. Furthermore, although JIA is often referred to as an “autoimmune” disease, we have shown that neutrophils from children with JIA show numerous abnormalities [18, 19] that do not correct even after children attain remission [20]. The finding that one of the JIA-associated genetic variants that we identified impacts enhancer function in both lymphoid and myeloid cells further supports our idea that neutrophil dysfunction is an essential feature of the disease rather than an epiphenomenon.

We note that we identified 2 genetic variants that altered enhancer function, one (rs117119468 C>T) in both Jurkat and HL60 cells, and the other (rs12722502 C>T) exclusively in Jurkat cells (**Figure 4**). This finding raises an interesting point that may need to be considered as we probe the other JIA risk loci and attempt to translate our understanding of genetics to improved patient care. It is possible, maybe likely, that none of the risk loci will have single causal variants, but, rather, multiple variants that may impinge on different functions to different degrees in different patients. Such a situation might explain, among other things, the considerable phenotypic variation that we see among children subsumed under the single disease entity, “polyarticular JIA.” So-called phenome-wide association studies[21] are likely to be a powerful tool that will allow us to link genetic variants with known/established biological effects with specific JIA phenotypes.

There are several limitations to the current study that must be acknowledged. The first is the nature of the assay used. The reporter plasmids may not accurately reflect the function of native chromatin [22], and results must therefore be placed in context with other information re: local chromatin structure. We have recently mined the abundant HiC data from multiple cell types available on public databases and identified physical contact between the first intron of the *IL2RA* gene and other regions on chromosome 10 (manuscript in preparation), so it is unlikely that a positive signal on luciferase assays represents experimental artifact. We also note that we have not identified the gene(s) regulated by this intronic enhancer. While intronic enhancers frequently regulate the genes in which they are located[23], they may also regulate other genes, typically within the same chromatin loop or toplologically-associated domain or TAD [24]. There are several genes of immunological interest within the TAD that encompasses the *IL2RA* haplotype, including the gene for IL15RA; IL15 and IL2 share both structural and functional homology [25] and it is plausible to hypothesize their expression/function may be regulated by common mechanisms.

In conclusion, we have demonstrated that the JIA-associated *IL2RA* locus has non-coding functions that can be altered by JIA-associated genetic variants. This work supports the concept that JIA causal variants are likely to disrupt regulatory functions rather than the coding sequence of the genes nearest the index SNPs identified on GWAS. This work also demonstrates that there are unlikely to be single causal variants operating at each risk locus, even in a single disease phenotype.

## Materials and Methods

### Cells and cell lines

Jurkat T cells and myeloid HL60 cells were obtained from the American Tissue Type Collection. We included myeloid HL60 cells in this study because the chromatin architecture in the region of interest is nearly identical in both CD4+ T cells and neutrophils. We confirmed the identity of these cell lines using short tandem repeat (STR) loci. STR markers are polymorphic DNA loci that contain repeated nucleotide sequences, and the number of repeats varies for each individual. Combinations of repeats were used to match cell lines to their original reported profile.

Jurkat T cells were grown and maintained in RPMI-1640 supplemented with penicillin G (100 U/ml), streptomycin (100 μg/ml) and 10% heat-inactivated FBS, while HL60 cells were grown and maintained in IMDM supplemented with penicillin G (100 U/ml), streptomycin (100 μg/ml) and 20% heat-inactivated FBS; and the cells were cultured at 37°C, 5% CO2 in a humidified atmosphere.

### Defining the IL2RA haplotype

We used the publically-available rAggr software program (http://raggr.usc.edu) to define the haplotype encompassing the SNP, rs7909519, the tag SNP used by Hinks et al [4] to identify *IL2RA* as a risk locus for JIA. We queried HG19 using r^2^ =0.8-1.0 and a minor allele frequency (MAF) of 0.001. This approach identified a 25,937 bp region spanning chr10:6,071,347-6,097,283 (hg19), as shown in **Figure 1**.

### Selection of regions to test

We used H3K4me1/H3K27ac ChIPseq data from Roadmap Epigenomics (accession numbers GSM1220567, GSM1220560 and GSM1102805) as well as our own H3K4me1/H3K27ac ChIPseq data in neutrophils (GSE66896)[8] to identify a 1500 bp region within the first intron of *IL2RA* that was likely to have functional activity (**Figure 2**). This region has prominent H3K4me1 and H3K27ac marks in both cell types as well as abundant transcription factor binding sites (ENCODE and Roadmap Epigenomics data). Subsequent studies were focused on 657bp (chr10:6092661-6093317) within this region.

### Purification and amplification of DNA by polymerase chain reaction (PCR)

DNA purification from whole blood was done using DNeasy Blood & Tissue Kit (Qiagen, USA). The purified DNA was used directly in the PCR. All PCR amplifications mixtures were carried out in a total volume of 50 μL containing 25 μL NEBNext® High-Fidelity 2X PCR Master Mix (New England BioLabs, Lpswich, MA, USA), 80 ng genomic DNA as a template, 50 pmol final concentrations from each of the forward and reverse primers, and the volume was completed to 50 μL using nuclease-free sterile water. The oligonucleotide sequences for cloning were FP: 5′- GTGGCCATGAGGAATCTGTAAG – 3′ and RP: 5′- CGACAATGAAACCAGGCAGA-3′ and the product size was 539bp (chr10:6092661-6093199); FP: 5′- CAGGTCCTCTTGCACTAGATTG-3′ and RP: 5′-AAGTTGACGTCAGCCTCTTC-3′ and the product size was 533bp (chr10:6092785- 6093317); FP: 5′- GAGGTGATGTGTTCTTCTCATCT-3′ and RP: 5′-TCATTTCTTCCTTGCTGTCCT-3′ and the product size was 402bp (chr10:6,089,866- 6,090,267). FP: 5′- GTGGCCATGAGGAATCTGTAAG-3′ and RP: 5′-AAGTTGACGTCAGCCTCTTC-3′ and the product size was 657bp (chr10:6092661- 6093317). XhoI (5′-C^TCGAG-3′) and BglII(5′-A^GATCT-3′) restriction endonuclease recognition sites were incorporated in the oligonucleotides for directional cloning.

### Cloning of the IL2RA Enhancer

The PCR amplified fragments were digested using XhoI and BglII (New England Biolabs, MA, USA) and ligated into a pGL4.23 vector (Promega E8411, Madison, MI, USA), previously digested using the same endonucleases. After an overnight incubation at 16°C, the ligation mixtures were transformed into competent *E. coli* JM109 cells (Promega). Plasmid DNA was isolated by Miniprep kit (Qiagen) according to the manufacturer’s instructions. All recombinant plasmid sequences were confirmed by Sanger sequencing.

### Site-Directed mutagenesis

Mutations were introduced with the QuikChange II Site-Directed Mutagenesis Kit (Agilent, Santa Clara, CA). The primers were designed according to the manufacturer’s protocol, utilizing the QuikChange Primer Design Tool (http://www.genomics.agilent.com). The wild-type (wt) construct pGL4.23-IL2RA enhancer (chr10:6092661-6093317) was used as template to generate 4 enhancer genetic variants (chr10:6092989, C>T (C239T); FP: AATAAAAGAGCCATCTCTCCAGACATCTGTCTGACG, RP: CGTCAGACAGATGTCTGGAGAGATGGCTCTTTTATT. chr10: 6093139, C>T(C479T); FP: GTGGTCTCTGCTTCCAGTCCTTGTGGTAGCA, RP: TGCTACCACAAGGACTGGAAGCAGAGACCAC. chr10: 6092686, A>G (A26G); FP:TGCGTGCTCTGCTTCTGCGATCTTACAGATTCCTC, RP: GAGGAATCTGTAAGATCGCAGAAGCAGAGCACGCA. chr10: 6093178, G>T (G518T); FP:GAAACCAGGCAGAGACTGCAACCCAGCTG and RP:CAGCTGGGTTGCAGTCTCTGCCTGGTTTC).

### Luciferase assays

Cells were transiently transfected using electroporation following the protocol of the manufacturer (Nucleofector Device with Cell line kit V, Lonza, USA). Cells (2×10^6^) were tranfected with 1ug of the pGL4.23-constructs plus 1 ug of the control Renilla plasmid DNA (pRL-TK; Promega) to normalize for transfection efficiency. Twenty-four hours after transfection, the cells were harvested, lysed and analyzed for luciferase activity. Luminescence was measured with the Dual-Luciferase® Reporter Assay System (Promega) in a microplate luminometer (Biotek Cytation 5MV Microplate Reader, Biotek). Luciferase activities are representative of at least three independent transfection experiments. Student’s t test was used to determine statistical significance

## Acknowledgements

This work has not been previously presented at a conference or published as a conference abstract.

## References

1. Prahalad S, O’Brien E, Fraser AM, Kerber RA, Mineau GP, Pratt D, et al. Familial aggregation of juvenile idiopathic arthritis. Arthritis Rheum. 2004;50(12):4022–7. Epub 2004/12/14. doi: 10.1002/art.20677. PubMed PMID: 15593218.

2. Hersh AO, Prahalad S. Immunogenetics of juvenile idiopathic arthritis: A comprehensive review. J Autoimmun. 2015;64:113–24. Epub 2015/08/26. doi: 10.1016/j.jaut.2015.08.002. PubMed PMID: 26305060; PubMed Central PMCID: PMCPMC4838197.

3. Herlin MP M.B.; and Herlin, T. Update on Genetic Susceptibility and Pathogenesis in Juvenile Idiopathic Arthritis. EMJ Eur Medical J Rheumatol. 2014;1.

4. Hinks A, Cobb J, Marion MC, Prahalad S, Sudman M, Bowes J, et al. Dense genotyping of immune-related disease regions identifies 14 new susceptibility loci for juvenile idiopathic arthritis. Nat Genet. 2013;45(6):664–9. Epub 2013/04/23. doi: 10.1038/ng.2614. PubMed PMID: 23603761; PubMed Central PMCID: PMCPMC3673707.

5. Cannon ME, Mohlke KL. Deciphering the Emerging Complexities of Molecular Mechanisms at GWAS Loci. Am J Hum Genet. 2018;103(5):637–53. Epub 2018/11/06. doi: 10.1016/j.ajhg.2018.10.001. PubMed PMID: 30388398; PubMed Central PMCID: PMCPMC6218604.

6. Boyle EA, Li YI, Pritchard JK. An Expanded View of Complex Traits: From Polygenic to Omnigenic. Cell. 2017;169(7):1177–86. Epub 2017/06/18. doi: 10.1016/j.cell.2017.05.038. PubMed PMID: 28622505; PubMed Central PMCID: PMCPMC5536862.

7. Wong L, Jiang K, Chen Y, Jarvis JN. Genetic insights into juvenile idiopathic arthritis derived from deep whole genome sequencing. Sci Rep. 2017;7(1):2657. Epub 2017/06/03. doi: 10.1038/s41598-017-02966-9. PubMed PMID: 28572608; PubMed Central PMCID: PMCPMC5453970.

8. Jiang K, Zhu L, Buck MJ, Chen Y, Carrier B, Liu T, et al. Disease-Associated Single-Nucleotide Polymorphisms From Noncoding Regions in Juvenile Idiopathic Arthritis Are Located Within or Adjacent to Functional Genomic Elements of Human Neutrophils and CD4+ T Cells. Arthritis Rheumatol. 2015;67(7):1966–77. Epub 2015/04/03. doi: 10.1002/art.39135. PubMed PMID: 25833190; PubMed Central PMCID: PMCPMC4485537.

9. Zhu L, Jiang K, Webber K, Wong L, Liu T, Chen Y, et al. Chromatin landscapes and genetic risk for juvenile idiopathic arthritis. Arthritis Res Ther. 2017;19(1):57. Epub 2017/03/16. doi: 10.1186/s13075-017-1260-x. PubMed PMID: 28288683; PubMed Central PMCID: PMCPMC5348874.

10. Hinks A, Ke X, Barton A, Eyre S, Bowes J, Worthington J, et al. Association of the IL2RA/CD25 gene with juvenile idiopathic arthritis. Arthritis Rheum. 2009;60(1):251–7. Epub 2009/01/01. doi: 10.1002/art.24187. PubMed PMID: 19116909; PubMed Central PMCID: PMCPMC2963023.

11. Pesenacker AM, Wedderburn LR. T regulatory cells in childhood arthritis--novel insights. Expert Rev Mol Med. 2013;15:e13. Epub 2013/12/04. doi: 10.1017/erm.2013.14. PubMed PMID: 24294966.

12. Knowlton N, Jiang K, Frank MB, Aggarwal A, Wallace C, McKee R, et al. The meaning of clinical remission in polyarticular juvenile idiopathic arthritis: gene expression profiling in peripheral blood mononuclear cells identifies distinct disease states. Arthritis Rheum. 2009;60(3):892–900. Epub 2009/02/28. doi: 10.1002/art.24298. PubMed PMID: 19248118; PubMed Central PMCID: PMCPMC2758237.

13. Jiang K, Wong L, Sawle AD, Frank MB, Chen Y, Wallace CA, et al. Whole blood expression profiling from the TREAT trial: insights for the pathogenesis of polyarticular juvenile idiopathic arthritis. Arthritis Res Ther. 2016;18(1):157. Epub 2016/07/09. doi: 10.1186/s13075-016-1059-1. PubMed PMID: 27388672; PubMed Central PMCID: PMCPMC4936089.

14. Hu Z, Jiang K, Frank MB, Chen Y, Jarvis JN. Complexity and Specificity of the Neutrophil Transcriptomes in Juvenile Idiopathic Arthritis. Sci Rep. 2016;6:27453. Epub 2016/06/09. doi: 10.1038/srep27453. PubMed PMID: 27271962; PubMed Central PMCID: PMCPMC4895221.

15. Wong L, Jiang K, Chen Y, Hennon T, Holmes L, Wallace CA, et al. Limits of Peripheral Blood Mononuclear Cells for Gene Expression-Based Biomarkers in Juvenile Idiopathic Arthritis. Sci Rep. 2016;6:29477. Epub 2016/07/08. doi: 10.1038/srep29477. PubMed PMID: 27385437; PubMed Central PMCID: PMCPMC4935846.

16. Farh KK, Marson A, Zhu J, Kleinewietfeld M, Housley WJ, Beik S, et al. Genetic and epigenetic fine mapping of causal autoimmune disease variants. Nature. 2015;518(7539):337–43. doi: 10.1038/nature13835. PubMed PMID: 25363779; PubMed Central PMCID: PMCPMC4336207.

17. Ernst J, Kheradpour P, Mikkelsen TS, Shoresh N, Ward LD, Epstein CB, et al. Mapping and analysis of chromatin state dynamics in nine human cell types. Nature. 2011;473(7345):43–9. doi: 10.1038/nature09906. PubMed PMID: 21441907; PubMed Central PMCID: PMCPMC3088773.

18. Jarvis JN, Petty HR, Tang Y, Frank MB, Tessier PA, Dozmorov I, et al. Evidence for chronic, peripheral activation of neutrophils in polyarticular juvenile rheumatoid arthritis. Arthritis Res Ther. 2006;8(5):R154. doi: 10.1186/ar2048. PubMed PMID: 17002793; PubMed Central PMCID: PMCPMC1779452.

19. Jarvis JN, Jiang K, Petty HR, Centola M. Neutrophils: the forgotten cell in JIA disease pathogenesis. Pediatr Rheumatol Online J. 2007;5:13. doi: 10.1186/1546-0096-5-13. PubMed PMID: 17567896; PubMed Central PMCID: PMCPMC1904449.

20. Jiang K, Frank M, Chen Y, Osban J, Jarvis JN. Genomic characterization of remission in juvenile idiopathic arthritis. Arthritis Res Ther. 2013;15(4):R100. doi: 10.1186/ar4280. PubMed PMID: 24000795; PubMed Central PMCID: PMCPMC4062846.

21. Bush WS, Oetjens MT, Crawford DC. Unravelling the human genome-phenome relationship using phenome-wide association studies. Nat Rev Genet. 2016;17(3):129–45. doi: 10.1038/nrg.2015.36. PubMed PMID: 26875678.

22. Inoue F, Kircher M, Martin B, Cooper GM, Witten DM, McManus MT, et al. A systematic comparison reveals substantial differences in chromosomal versus episomal encoding of enhancer activity. Genome Res. 2017;27(1):38–52. doi: 10.1101/gr.212092.116. PubMed PMID: 27831498; PubMed Central PMCID: PMCPMC5204343.

23. Shaul O. How introns enhance gene expression. Int J Biochem Cell Biol. 2017;91(Pt B):145–55. doi: 10.1016/j.biocel.2017.06.016. PubMed PMID: 28673892.

24. Krijger PH, de Laat W. Regulation of disease-associated gene expression in the 3D genome. Nat Rev Mol Cell Biol. 2016;17(12):771–82. doi: 10.1038/nrm.2016.138. PubMed PMID: 27826147.

25. Bodnar A, Nizsaloczki E, Mocsar G, Szaloki N, Waldmann TA, Damjanovich S, et al. A biophysical approach to IL-2 and IL-15 receptor function: localization, conformation and interactions. Immunol Lett. 2008;116(2):117–25. doi: 10.1016/j.imlet.2007.12.014. PubMed PMID: 18280585.

